# Axo-vascular coupling mediated by oligodendrocytes

**DOI:** 10.1101/2022.06.16.495900

**Authors:** Alejandro Restrepo, Andrea Trevisiol, Camilo Restrepo-Arango, Constanze Depp, Andrew Octavian Sasmita, Annika Keller, Iva D. Tzvetanova, Johannes Hirrlinger, Klaus-Armin Nave

## Abstract

The high energy requirements of the cortical gray matter are met by the precise cooperation of neurons, glia, and vascular cells in a process known as neurovascular coupling (NVC). In contrast, the existence and significance of NVC in white matter (WM) are still debated and basic regulatory mechanisms are unknown. We recently discovered that oligodendrocytes sense the spiking axons’ activity via NMDA receptors and regulate their cell surface expression of glucose transporter GLUT1 allowing an increase in glycolytic metabolism that enables lactate release to metabolically support the axons. Here, we show for the mouse optic nerve (ON), a model WM tract, that the vascular support is also dynamically controlled. Axonal spiking activity induces small vessel dilations which are sustained for more than 20 minutes upon the ending of electrical stimulation. Pharmacological inhibition shows that the electrically evoked dilation is mediated by the prostaglandin E_2_ receptor EP_4_ and can be modulated by the oxygen concentration, as has been shown in the grey matter. Importantly, we found in ONs from conditional mouse mutants that oligodendroglial NMDA receptors are required for this type of neurovascular response, demonstrating a critical role of oligodendrocytes in coupling axonal activity to pericyte function. Reminiscent of NVC in cortical slices, the “axo-vascular” response is slower and may represent a more rudimentary form of neurovascular coupling.

## Introduction

The cerebral WM in adult brains is particularly vulnerable to cerebrovascular diseases such as ischemia and to vascular risk factors like hypertension (Longstreth et al., 1996; Pantoni et al., 1996). Loss of cerebrovascular function and microcirculatory failures are important contributors to white matter abnormalities (Joutel et al., 2010; Prins and Scheltens, 2015). Thus, the normal WM homeostasis and axonal functions are highly dependent on the vascular system, which supplies metabolites and oxygen for ATP production. Under healthy conditions, these metabolites are almost exclusively glucose and lactate (Clarke and Sokoloff, 1999), and on rare occasions ketone bodies (Stumpf et al., 2019).

In grey matter, synaptic communication is the most energy-consuming process (Howarth et al., 2012; Yu et al., 2018). Synaptic activity leads to a metabolic shift in astrocytes and neurons which increases glucose uptake, lactate production, and a metabolite flux between these two cells to support synaptic ATP production (Bak et al., 2009; Chuquet et al., 2010; Díaz-García et al., 2021; Dienel, 2012; Kasischke et al., 2004; Kasparov, 2016; Nehlig et al., 2004; Rouach et al., 2008; Yellen, 2018). Ultimately, the metabolite flux is supported by the vascular system by providing glucose and oxygen. By several proposed mechanisms (Hosford and Gourine, 2019), blood vessels dilate and regional cerebral blood flow (CBF) increases upon synaptic activity, most prominently via astroglial release of vasoactive prostaglandins (Hall et al., 2014; Mishra et al., 2016), lactate (Gordon et al., 2008), or potassium ions (Glück et al., 2021; Longden et al., 2017). The CBF peak is preceded by a decrease in both glucose and oxygen concentration (Lecoq et al., 2011; Wei et al., 2016). The increased flow produces a higher increase in oxygen availability than its metabolism (Lecoq et al., 2009; Parpaleix et al., 2013) which is necessary to maintain normal oxygen levels in regions away from the local stimulation (Devor et al., 2011). Disruptions in NVC contribute to pathologies observed in neurodegenerative diseases (Iadecola, 2013; Kisler et al., 2017; Montagne et al., 2018).

Despite WM using on average less energy than grey matter, upon electrical stimulation, there is an increased rate of glucose consumption in WM (Weber et al., 2002), suggesting WM requires a modulated delivery of glucose and oxygen, possibly through similar NVC mechanisms to those observed in grey matter. Because spiking axons cannot cover their ATP consumption by glycolysis and they rely on the support of oligodendrocytes and astrocytes (Lee et al., 2012; Meyer et al., 2018; Philippot et al., 2021; Trevisiol et al., 2020), oligodendrocytes need to be “aware” of axonal activity. In fact, it is thought that axonal spiking activity causes a minor non-synaptic release of glutamate which can be picked up by oligodendrocytes via NMDARs that are facing the periaxonal space (Doyle et al., 2018; Kukley et al., 2007; Micu et al., 2016). Indeed, NMDAR activation on oligodendrocytes triggers relocalization of GLUT1 that leads to an increased glucose flux into the cell that increases oligodendroglial glycolysis and production of lactate. Lactate is shuttled back to axons via monocarboxylate transporters to support axonal ATP synthesis (Saab et al., 2016). In fact, axonal ATP levels are more dependent on glial lactate production than glucose availability (Trevisiol et al., 2017). However, to fully exploit this lactate shuttle, oxygen must not be the limiting factor during the oxidation of lactate in axonal mitochondria, suggesting that WM may have a higher need for oxygen than glucose.

Given that WM has an activity-dependent glucose metabolism, that oligodendrocytes support axons through lactate following NMDAR activation, and that oxygen is critical in WM metabolism, we hypothesize that an efficient metabolic support of axons by oligodendrocytes ultimately also requires the participation of the vascular system. To address the question “how is neurovascular coupling being regulated in a WM tract?”, we used the well-studied *ex vivo* optic nerve (ON) preparation (Stys et al., 1991). Our data support the existence of an “axo-vascular” coupling in WM and, similar to what has been previously described for the grey matter, this coupling mechanism appears to be modulated by the concentration of oxygen and mediated by the prostaglandin E_2_ receptor EP_4_. Moreover, we suggest a working model, in which oligodendroglial NMDARs detect axonal spiking activity to turn this into a functional axo-vascular response.

## Results

### The ON is a highly vascularized tissue

In the absence of prior studies devoted to NVC in mammalian WM tracts, we focused on a well-known model system, the rodent optic nerve (ON). Using confocal and light-sheet microscopy with a modified iDisco clearing protocol (Depp et al., 2021), we revealed the 3D vessel-arrangement of the ON and annotated different populations of perivascular cells based on a recently proposed nomenclature for their morphology (Grant et al., 2019).

The adult mouse ON receives input from one arteriole, localized in a caudal position close to the optic chiasm (Figure 1A, dashed white arrows, and Figure 1C). This arteriole branches into several pre-capillary arterioles (Figure 1C). One of them runs almost completely to the rostral end of the nerve (Figure 1A, white arrowheads), at a depth of 10 µm – 40 µm from the surface of the nerve (Figure 1B). Our results show that the arteriole runs inside the nerve, i.e. within the first 27% of the total nerve’s depth, as opposed to outside of the fiber bundle as suggested previously (Barbosa et al., 2022; May and Lütjen-Drecoll, 2002). A vein several times the size of the arteriole, runs at a depth of 65 µm – 90µm along the entire nerve’s length (Figure 1A, dashed yellow arrows), reaching the most caudal part, where it branches into several venules (Figure 1A, yellow arrows).

**Figure 1:**
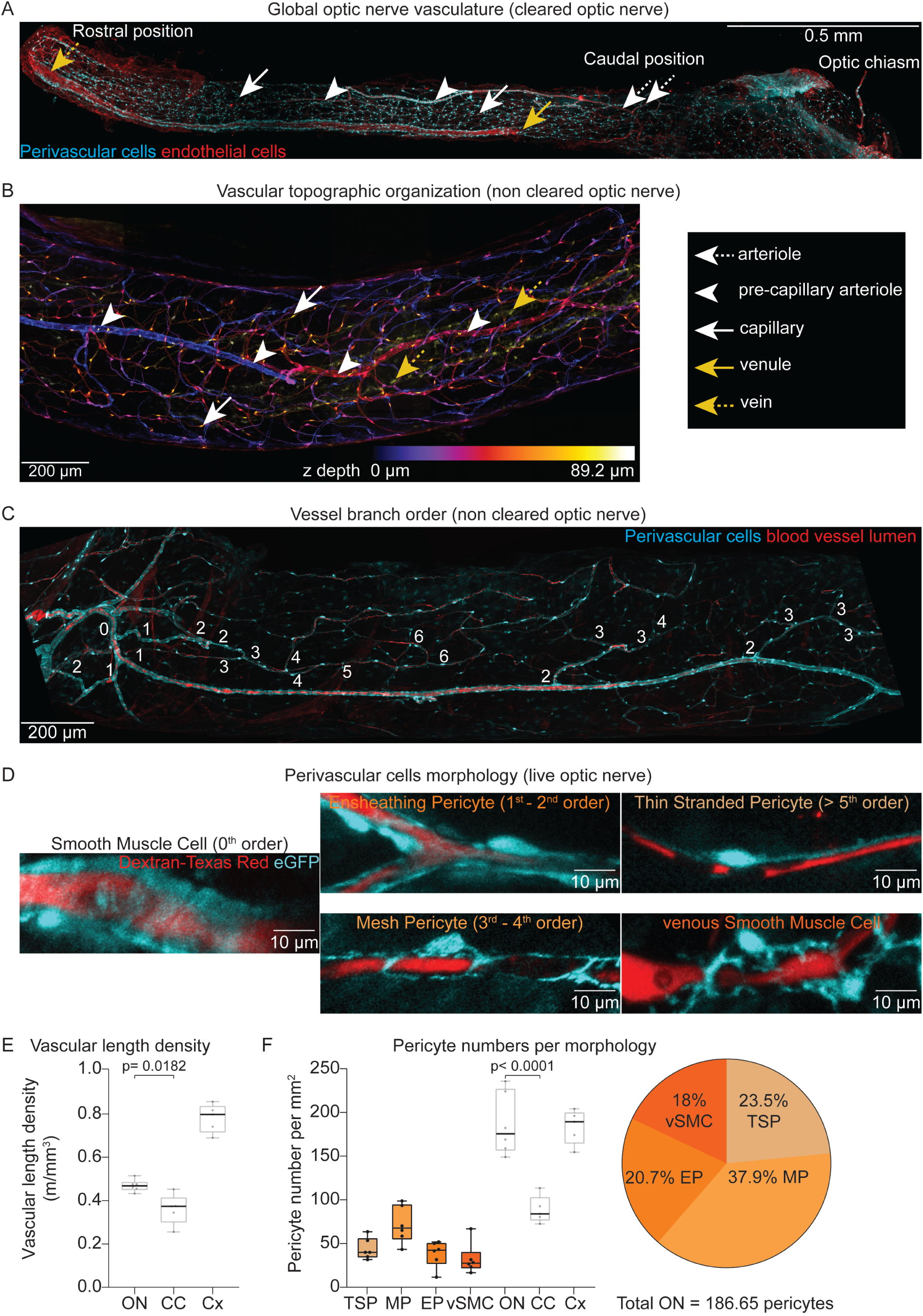
The optic nerve is highly vascularized. **A)** Representative light-sheet microscopy image of a cleared ON. Perivascular cells express cytosolic eGFP (cyan) and the endothelial cells were labeled by staining for podocalyxin (red). **B)** Representative image showing the difference in vessel type depending on the depth. Note how the pre-capillary arteriole (white arrowheads) are on the opposite side of the veins (yellow dashed arrows). **C)** Branching order of the ON vasculature. **D)** Representative images of different perivascular cell morphology in the ON. **E)** Quantification of the vascular length density of the ON, corpus callosum (CC), and cortex (Cx). **F)** Quantification of pericyte numbers per morphology in the ON. The box plots on the left show the number of pericytes averaged across 4 images per ON in 6 different mice (n= 6). TSO: thin-stranded pericytes, MP: mesh pericytes, EP: ensheathing pericytes, vSMC: venous smooth muscle cell. The total number of pericytes in the ON is also reported (white box plots). For comparison, the total number of pericytes in the corpus callosum (CC) and cortex (Cx) are also shown. The pie chart on the right shows the relative percentage of pericyte types. In box plots, the central line shows the median, the edges of the box define the upper and lower quartile values, and whiskers show the minimum-maximum range.

To further characterize the vasculature of the nerve, we determined the morphology of the perivascular cells in relation to the vessel order (Figure 1C), the single arteriole being classified as order 0. Vessels of orders 1 and 2 are covered by ensheathing pericytes (Figure 1D). Vessels of orders 3 and 4 are covered by mesh pericytes (Figure 1D). The rest of the vessels (5 or higher), are covered by thin-stranded pericytes (Figure 1D). The venules or veins of the ON are covered by stellate pericytes or venous smooth muscle cells.

Strikingly, ONs are highly vascularized (Figure 1A, white arrows) with an average vascular length density of 0.47 m/mm^3^ ± 0.027 m/mm^3^, significantly higher than the corpus callosum (0.36 m/mm^3^ ± 0.071 m/mm^3^ – Figure 1E). This difference is also reflected by an increase in pericyte density. The ON exhibited 186.6 pc/mm^2^ ± 35.7 pc/mm^2^, about twice the number of pericytes observed in the corpus callosum (88.8 pc/mm^2^ ± 15.7 pc/mm^2^ – Figure 1F). Similar to hippocampus or visual cortex (Shaw et al., 2021), the majority of pericytes are mesh pericytes (71.7 pc/mm^2^ ± 21.1 pc/mm^2^; 37.9%), followed by thin-stranded pericytes (44.1 pc/mm^2^ ± 12.1 pc/mm^2^; 23.5%) and ensheathing pericytes (38.5 pc/mm^2^ ± 15 pc/mm^2^; 20.7%). Venous smooth muscle cells represent the smallest population with 32.4 pc/mm^2^ ± 17.7 pc/mm^2^ or 18%.

These images show that the vascularization of the ON is surprisingly high compared to the corpus callosum, but also, that the vascular organization of the ON is similar to that of grey matter.

### WM pericytes retain their contractile properties *ex vivo*

Next, we asked whether perivascular cells in the acutely excised ON would retain their contractile properties, allowing *ex vivo* studies on axo-vascular coupling (AVC). In cortical brain slices, NVC is studied by stimulating neurons of the cortex by eliciting field potentials (Bojovic et al., 2022; Mishra et al., 2014). Here, we used suction electrodes to evoke action potential propagation along most of the ON axons while simultaneously measuring the diameter of the vessels sitting 10 µm – 30 µm below the nerve’s surface using confocal microscopy (Figure 2A).

**Figure 2:**
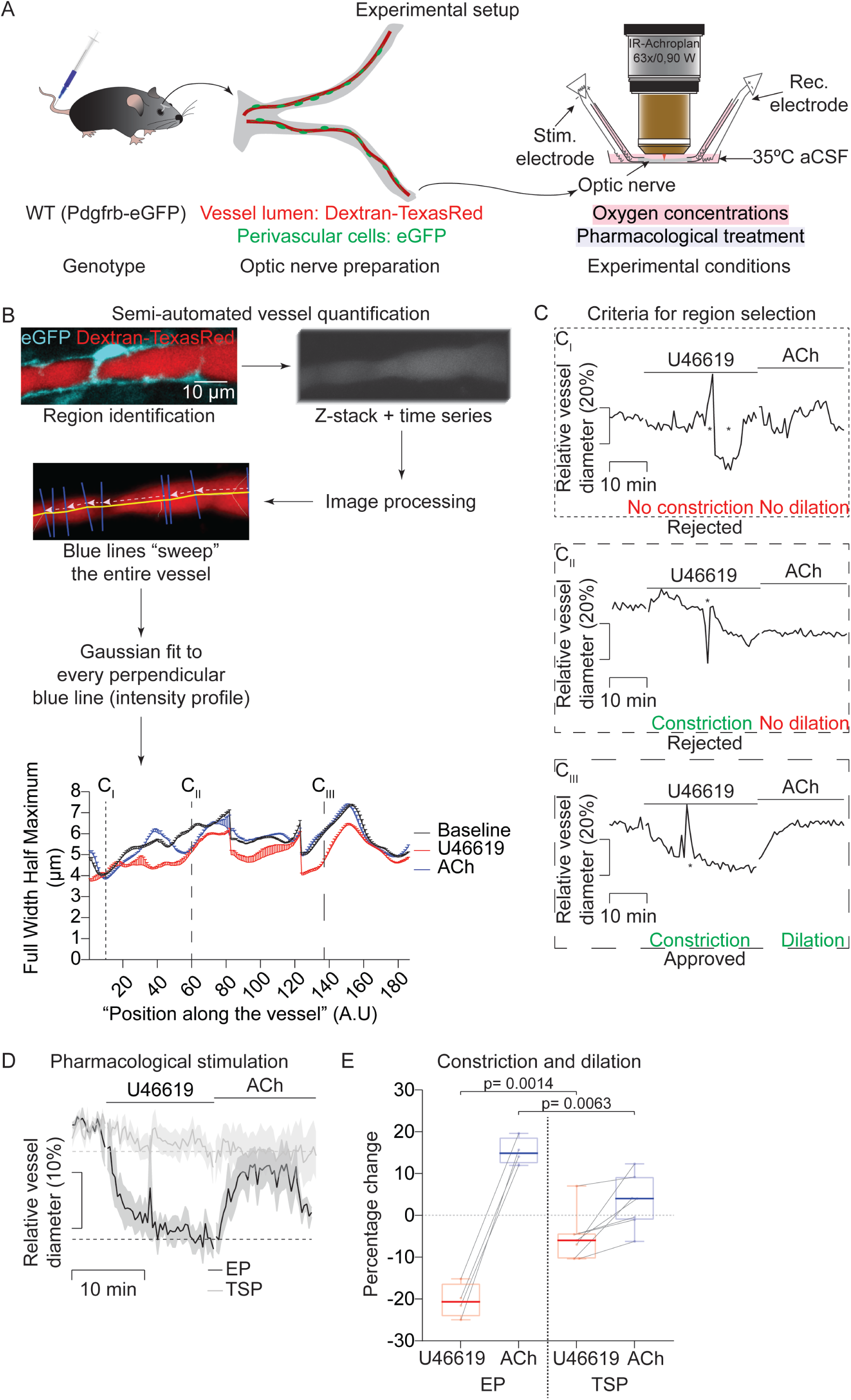
Vessel diameter quantification and perivascular cells contractility ex vivo. **A)** Diagram of the experimental setup used for live imaging and electrophysiological stimulation of the ON. Dextran-TexasRed is injected via tail-vein injection before ONs are dissected. One ON is clamped between two suction electrodes that allow stimulation and recording of the compound action potential of the nerve. Simultaneously, blood vessels, pericytes, or smooth muscle cells are imaged by confocal microscopy. **B)** Image analysis routine to determine the intensity profile along the blue lines perpendicular to the entire vessel (yellow line). The blue line moves along the entire vessel and calculates the intensity profile. Each intensity profile is fitted to a Gaussian curve and the Full-Width Half Maximum (FWHM) is calculated for each one as a proxy for vessel diameter. The FWHM of each line corresponding to the last 2 min of each treatment is plotted versus the position of the vessel. For clarity, data are presented as mean + standard error of the mean. **C)** Selection criteria for region selection for vessel diameter analysis. C_I_) Vessel diameter over the time at the position shown in B. There is no change in diameter with the different treatments. C_II_) There is a constriction but there is no dilation with ACh. C_III_) This position meets the two necessary criteria for analysis (treatment-induced constriction and dilation). The peaks marked with an asterisk during the U46619 treatment are due to a cell, possibly a monocyte, passing by inside the blood vessel during the acquisition. **D)** ONs were bathed with the vasoconstrictor U46619 (100 nM) for 15 min followed by 15 min application of acetylcholine (ACh – 100 µM), a vasodilator. Only the vessels with ensheathing pericytes (EP; n= 4) show a robust constriction and dilation upon vasoactive treatment. Vessels with thin-stranded pericytes (TSP; n= 7) show a very slow and small constriction but almost no dilation. The dotted line shows the vessel size at the end of the constriction. **E)** Quantification of the last 2 mins of constriction for both cell types and 2 mins at the maximal dilation during ACh treatment for both groups. The maximal dilation is defined as the point with the highest vessel dilation in the 15 min of ACh treatment. The difference between U46619 in EP and TSP and ACh in EP and TSP was calculated with an unpaired two-tailed t-test. In box plots, the central line shows the median, the edges of the box define the upper and lower quartile values, and the whiskers show the minimum-maximum range.

Contractile properties of pericytes have been debated (Hartmann et al., 2022). To evaluate if pericytes in the *ex vivo* ON have and retain their contractile properties, we applied the vasoconstrictor U46619 (100 nM) for 15 min followed by acetylcholine (ACh – 100 µM) as a vasodilator for 15 min, and evaluated subsequent changes in the vessel diameter (Figure 2D). Our data confirm that ensheathing pericytes (EPs) have robust contractile properties. EPs contract following U46619 treatment and cause the underlying vessel to constrict by 20.4% ± 3.5%, compared to baseline. These cells also respond to ACh application, causing the vessel to dilate by 15.0% ± 2.8%, over the constricted level (Figure 2E). In contrast, vessels with thin-stranded pericytes (TSP) constrict only by 5.1% ± 5.4%, i.e. 15.3% less than EPs (n EP= 4, n TSP= 7; p= 0.0014 using an unpaired two-tailed t-test). Similarly, vessels with TSPs dilate by 3.1% ± 5.8% over the constricted level, i.e. 12.2% less than Eps following ACh-induced dilation (n EP= 4, n TSP= 7; p= 0.0063 using an unpaired two-tailed t-test) (Figure 2E). These results confirm previous observations on the contractility of different pericyte types (Gonzales et al., 2020; Hartmann et al., 2021).

### Stimulation evoked dilation in the optic nerve

Next, we asked whether the electrical stimulation of the ON would elicit a vascular response, i.e. if NVC (or “axo-vascular coupling; AVC) is also present in a WM tract. As EPs show robust contractile properties in the *ex vivo* ON preparation upon vasoactive substance treatment, we only focused on these cells for the rest of the experiments.

We stimulated the nerve with 4 different frequencies: a baseline stimulation (0.1 Hz) (Figure 3B) caused no dilation (0.2% ± 1.0%) at the end of the 3 min stimulation (Figure 3G). The 4 Hz stimulation (Figure 3C) increased the vessel diameter by 1.2% ± 1.2% (Figure 3G). The 25 Hz stimulation (Figure 3D), elicited a 2.4% ± 1.8% increase in the vessel diameter (Figure 3G). Finally, the 100 Hz stimulation (Figure 3E) also induced an increase of 2.3% ± 1.6% in the vessel diameter (Figure 3G).

**Figure 3:**
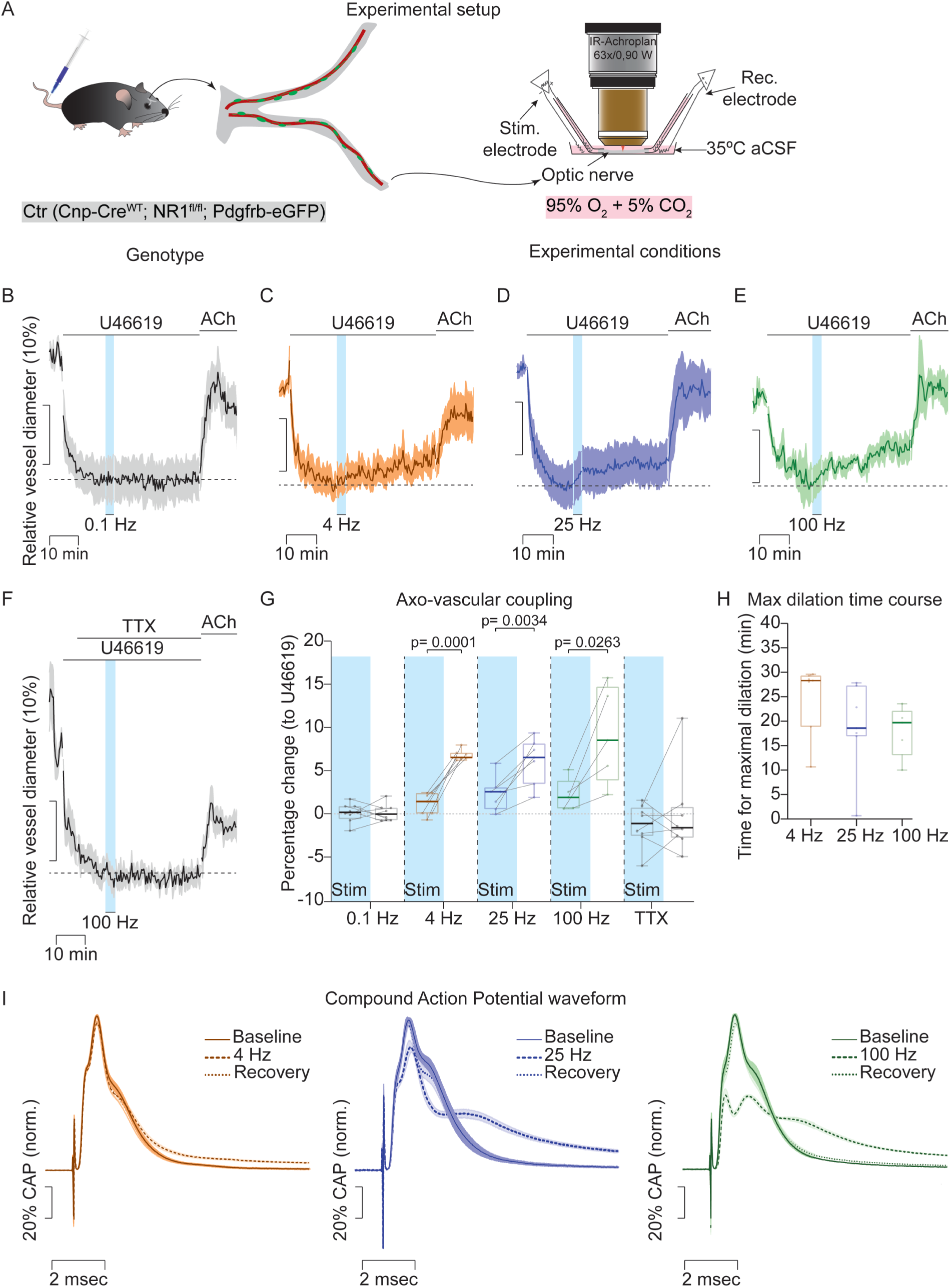
Stimulation evoked dilation in the ON. **A)** Diagram of the experimental setup used for live imaging and electrophysiological stimulation of the ON. The aCSF bath solution was oxygenated with carbogen and kept at 35°C. **B)** ONs were preconstricted with U46619 (100 nM) and stimulated at 0.1 Hz (n= 8), **C)** 4 Hz (n= 7), **D)** 25 Hz (n= 7), **E)** 100 Hz (n= 5), and **F)** 100 Hz under TTX (1 µM; n= 8) for 3 min followed by a post-stimulation period of 30 min. The dotted line shows the preconstriction vessel diameter. The blue shaded area indicates the stimulation period. **G)** Quantification of the dilation (percentage) of the vessels during the stimulation (blue shaded area) and the post-stimulation period. **H)** Time for vessels to reach the maximal dilation during the post-stimulation period of 4 Hz, 25 Hz, and 100 Hz. **I)** Compound action potential waveform of ONs during 4 Hz, 25 Hz, and 100 Hz stimulations. Plotted is the average waveform at the end of the baseline (solid line), at the end of the stimulation (dashed line), and after 5 min of post-stimulation (dotted line). In box plots, the central line shows the median, the edges of the box define the upper and lower quartile values, and the whiskers show the minimum-maximum range.

Unexpectedly, vessel dilation slowly progressed during the following 30 min of post-stimulation, rather than falling back to baseline as observed in grey matter. Vessels reached a maximal dilation during this period (Figure 3C-E). At the end of the 30 min of 4 Hz post-stimulation, vessels dilated a further 5.6% and reached a maximum vessel diameter of 6.9% ± 0.5% (n= 7; p= 0.0001 compared to stimulation (1.2% ± 1.2%); paired two-tailed t-test). This same response was observed during the post-stimulation period of the 25 Hz (6.2% ± 2.4%, (n= 7; p= 0.0034 compared to stimulation (2.4% ± 1.8%); paired two-tailed t-test) and 100 Hz (9.2% ± 5.0%, (n= 5; p= 0.0263 compared to stimulation (2.3% ± 1.6%); paired two-tailed t-test) stimulations (Figure 3G). In contrast, the baseline stimulation did not lead to sustained dilation (0.3% ± 0.9%, n= 8). To confirm that the axonal activity was responsible for the observed vessel dilation, we incubated the ONs in tetrodotoxin (TTX – 1 μM) before the onset of stimulation (Figure 3F). TTX effectively blocked the stimulation-evoked vessel dilation and the delayed dilation during the post-stimulation period, (−1.2% ± 2.3% and −0.06% ± 4.5%, n= 8, respectively), confirming that axonal activity drives vasodilation in the ON (Figure 3G).

To further characterize the kinetics of the ON’s AVC response, we determined the time courses necessary to reach the maximal dilation *ex vivo*. For the 25 Hz stimulation, we calculated 18.9 min ± 8.5 min for the vessels to reach their maximal dilation. Interestingly, no difference was observed between 4 Hz, 25 Hz, and 100 Hz (24.8 min ± 6.7 min for 4 Hz; 18.9 min ± 8.5 min for 25 Hz; 18.0 min ± 4.6 min for 100 Hz, respectively), with the following non-significant p-values (comparison 4 Hz-25 Hz): 0.1920; (comparison 4 Hz-100 Hz): 0.1660; (comparison 25 Hz-100 Hz): >0.99 using a Kruskal-Wallis test with multiple comparisons using 3 pairs (Figure 3H). As discussed below, these response times are distinct from the rapid NVC at cortical synapses (Mishra et al., 2016).

To evaluate if the chosen stimulation paradigm had caused an ON injury that could be responsible for the sustained dilation, we qualitatively compared the compound action potential (CAP) for each stimulation frequency (Figure 3I). The waveform of the CAPs showed complete recovery at each frequency, indicating that the conduction properties of the acutely isolated nerves were not impaired by the stimulation paradigm.

### Oxygen modulates the AVC response in the optic nerve

Different mechanisms have been proposed for grey matter NVC (Hosford and Gourine, 2019). A widely held view holds that astrocytes use COX1 to produce prostaglandin E_2_ (PGE_2_) which is secreted to activate the EP_4_ receptor on perivascular cells, inducing vasodilation (Gordon et al., 2008; Mishra et al., 2016). This NVC response is modulated by the oxygen concentration in *ex vivo* experiments (Gordon et al., 2008; Hall et al., 2014; Mishra et al., 2011). To explore whether PGE_2_-EP_4_ receptor signaling is involved in WM AVC, we used pharmacological inhibition of the EP_4_ receptor at different oxygen concentrations and evaluated the vessel dilation as a function of electrical stimulation (Figure 4A).

**Figure 4:**
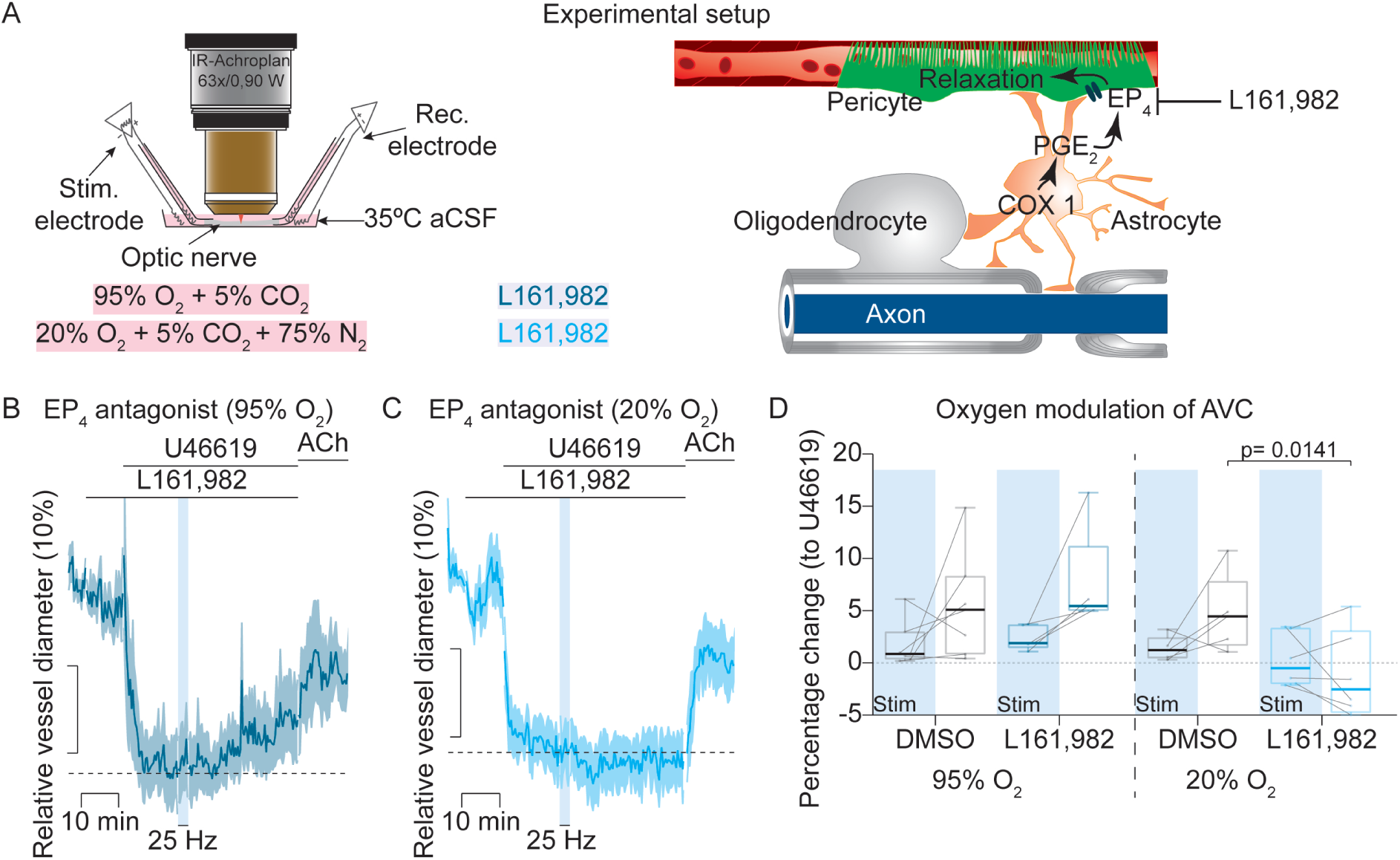
Oxygen concentration modulates the NVC response in the ON. **A)** Diagram of the experimental setup used for live imaging and electrophysiological stimulation of the ON. The aCSF bath solution was oxygenated with carbogen (95% O2 + 5% CO2) or with a mixture of 20% O2 + 5% CO2 + 75% N_2_, and the EP4 receptor antagonist L161,982 was used. **B)** Relative vessel diameter in ONs treated with the EP_4_ receptor antagonist at 95% oxygen and **C)** at 20% oxygen. The dotted line shows the preconstriction vessel diameter. The blue shaded area shows the stimulation window. **D)** Quantification of the relative vessel diameter after stimulation at 25 Hz, comparing different oxygen concentrations. In box plots, the central line shows the median, the edges of the box define the upper and lower quartile values, and the whiskers show the minimum-maximum range.

At 95% oxygen (most frequently used by electrophysiologists), the EP_4_ receptor is not detectably involved in AVC (Figure 4B). Vessels of the nerves treated with the EP_4_ antagonist L161,982 (1 µM) dilated 7.6% ± 4.9% after 30 min post-stimulation. In control nerves, vessels dilated 5.4% ± 5.0% over the same period of time (p= 0.5948 with an unpaired Kruskal-Wallis test using 2 pairs for multiple comparisons; Figure 4E). These data indicate that AVC is not dependent on PGE_2_-EP_4_ signaling at 95% oxygenation.

At lower oxygen concentrations (20%), the relevant signaling pathways differ and EP_4_ receptor inhibition blocks the NVC response upon nerve stimulation (Figure 4C). Vessels of control nerves dilated 4.7% ± 3.7%, while L161,982 treatment induced a further constriction of 1.4% ± 4.9% over the preconstricted level (p= 0.0141, using an unpaired One-way ANOVA with multiple comparisons using 2 pairs) (Figure 4D, right). These data show that the oxygen concentration modulates the AVC response in WM, in a similar way as shown in grey matter.

### Oligodendroglial NMDA receptors participate in AVC

We have previously proposed a model of oligodendrocytes that modulate lactate release as a function of NMDA type glutamate receptor signaling, according to the electric activity of axons hey ensheath (Micu et al., 2018; Saab et al., 2016). Oligodendrocytes are thus in the position to “measure” the spiking-dependent axonal energy needs. We, therefore, hypothesized that oligodendroglial NMDARs are required for the establishment of AVC with oligodendrocytes being a “relay station” between axonal energy demand and vasodilation. By taking a loss-of-function approach, we studied AVC in oligodendrocyte-specific NR1 knockout ONs (termed cKO in the following; Figure 5).

**Figure 5:**
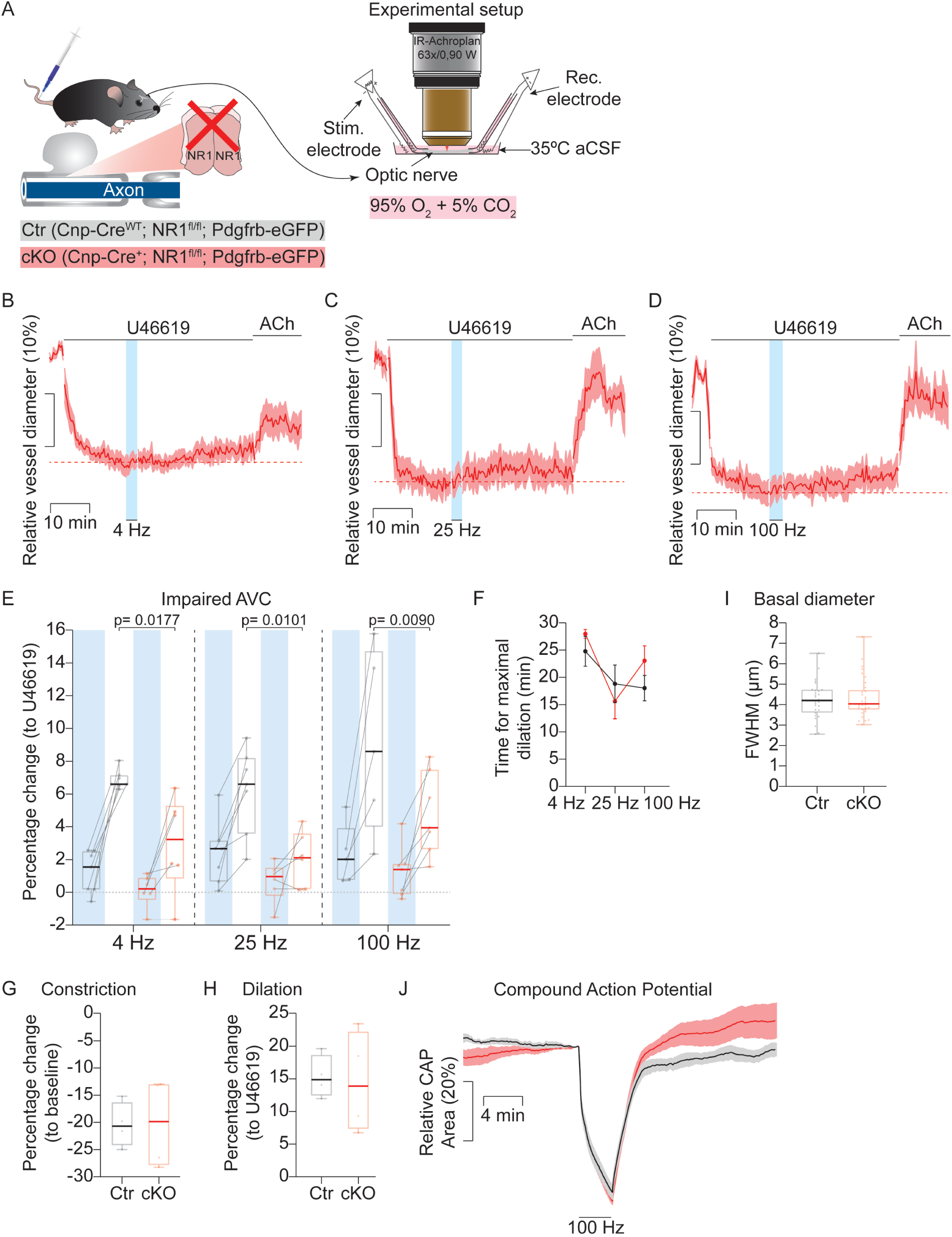
Oligodendroglial NMDA receptors are involved in WM NVC. **A)** Diagram of the experimental setup used for live imaging and electrophysiological stimulation of the ON. To test if oligodendroglial NMDA receptors are involved in NVC we used a previously described oligodendrocyte-specific NMDA receptor knock-out. **B)** ON from cKO mice were preconstricted with U46619 (100 nM) and stimulated at 4 Hz (n= 6), **C)** 25 Hz (n= 6), and **D)** 100 Hz (n= 7) for 3 min followed by a recovery period of 30 min. The dotted line shows the preconstriction level. The blue shaded area shows the stimulation period. **E)** Quantification of the dilation percentage of the vessels during the stimulation (blue shaded area) and the post-stimulation period (right) for each frequency of stimulation. At 4 Hz, vessels dilate 2.9% ± 2.7% (p= 0.0177, compared to Ctr; using an unpaired one-way ANOVA with 6 pairs for multiple comparisons); at 100 Hz vessels dilate 4.8 % ± 2.3 % (p= 0.0090, compared to Ctr; using an unpaired one-way ANOVA with 6 pairs for multiple comparisons). **F)** The time to reach the maximal dilation during the recovery period does not change with the deletion of NMDA receptors in oligodendrocytes. Upon 4 Hz stimulation cKO mice need 28 min ± 1.8 min to reach a maximal dilation (p> 0.99 using a Kruskal-Wallis test with 3 pairs of multiple comparisons). Upon 25 Hz stimulation, vessels of cKO mice dilated maximally at 15.6 min ± 7.2 min (p> 0.99 using a Kruskal-Wallis test with 3 pairs of multiple comparisons). After 100 Hz stimulation, vessels of cKO mice dilated maximally after 23.0 min ± 6.6 min (p= 0.5589 using a Kruskal-Wallis test with 3 pairs of multiple comparisons). **G)** Quantification of the last 2 min of preconstruction for both control animals (n= 4) and cKO animals (n= 4). Vessels constricted 20.2% ± 7.1% (p= 0.9702, compared to Ctr animals; using an unpaired two-tailed t-test). **H)** Quantification of 2 min located around the maximal dilation during ACh treatment for both groups. The maximal dilation is defined as the point with the highest vessel dilation in the 15 min of ACh treatment. Vessels dilated 15.3% ± 2.8% (p= 0.8499, compared to Ctr animals; using an unpaired two-tailed t-test). **I)** Vessel diameter is not changed in ex vivo ONs after oligodendroglial NMDA receptors deletion. Ctr: 4.2 µm ± 0.9 µm; cKO: 4.3 µm ± 0.9 µm; n Ctr = 28 and n cKO = 34; p= 0.9384 using a Mann-Whitney two-tailed test) **J)** Quantification of the area under the curve of CAP for 100 Hz for Ctr and cKO mice (n Ctr 100 Hz= 5; n cKO 100 Hz= 5; p= 0.31 using an unpaired two-tailed t-test). In box plots, the central line shows the median, the edges of the box define the upper and lower quartile values, and the whiskers show the minimum-maximum range.

The electrophysiological characterization of the cKO ONs revealed that AVC is impaired, regardless of the stimulation frequency (Figure 5B-D). For all frequencies, we observed a reduction of ∼1.2% of the vessel diameter during the stimulation period compared to control (Ctr) animals, shown in figure 3. Moreover, at 30 min post-stimulation the vessels from cKO animals also showed a sustained and increased dilation, but with a lower amplitude (−4.2% on average) compared to control mice. At 25 Hz the dilation was 2.0 % ± 1.5 % (p= 0.0101, compared to Ctr; using an unpaired one-way ANOVA with 6 pairs for multiple comparisons) (Figure 5E). Interestingly, the time needed to reach the maximal dilation in cKO was comparable to control ON (Figure 5F).

The smaller dilation observed in cKO nerves is unlikely caused by an intrinsic inability of vascular cells to respond because the overall degree of constriction and dilation following U46619 and ACh administration was unimpaired (constriction: p= 0.97, compared to controls; dilation: p= 0.85, compared to controls; using unpaired two-tailed t-tests) (Figure 5G and H). There were also no differences in basal vessel size in the ONs (Figure 5I) (Ctr: 4.22 µm ± 0.87 µm; cKO: 4.32 µm ± 0.93 µm; n Ctr = 28 and n cKO = 34; p=b0.94 using a Mann-Whitney two-tailed test). These data indicate that the impaired AVC seen in cKO mice is due to the lack of NMDARs in oligodendrocytes and is not caused by changes in vascular cell functionality.

To further confirm that the impaired NVC in cKO mice was not due to changes in basic axonal firing capabilities, we quantified the CAP of the nerves by calculating the area underneath the curve of the waveform of the CAP, which correlates with the number of firing axons in the tissue (Stys et al., 1991). Here, the CAP amplitudes were not different for the genotypes (100 Hz: p= 0.32; using an unpaired two-tailed t-test). This suggests that AVC impairment in cKO ONs was not caused by axonal firing defects (Figure 5J).

## Discussion

Although some functions of NVC in grey matter are still under investigation (Howarth et al., 2021), it is widely accepted that one role of NVC is to increase the local oxygen concentration according to the energy demands of electrically active brain regions (Devor et al., 2011). While it has been shown that WM is metabolically less active than gray matter (Attwell and Laughlin, 2001; Clarke and Sokoloff, 1999; Howarth et al., 2012), axons in the WM are shielded from the diffusion of energy substrates by myelin (Hirrlinger and Nave, 2014) and rely almost exclusively on oxidative glial lactate metabolism to produce their ATP (Trevisiol et al., 2017). This high axonal dependency on glucose and oxygen is also supported by the evidence of increased susceptibility of WM damage to ischemic events (Li et al., 2019).

For the cortical and hippocampal grey matter, it has been shown that an increase in neuronal activity prompts local blood vessels to increase their diameter, temporally providing an elevated supply of glucose and oxygen to match energy demands. When the higher spiking activity comes to an end, the vessel diameters fall back to baseline, as shown in several *in vivo* and in *ex vivo* studies (Chow et al., 2020; Hall et al., 2014; Mishra et al., 2016; Rungta et al., 2021; Zonta et al., 2003). This vascular response has been repeatedly shown to be modulated by at least 12 different receptors/molecules/enzymes (Hosford and Gourine, 2019). However, there is practically no information about vascular coupling in the WM, presumably because the corresponding readout is much lower, temporally delayed, or therefore simply missed. To the best of our knowledge, we have analyzed for the first time a vascular response to neuronal (axonal) spiking activity in a WM tract, the isolated ON. To our surprise, this response, which we termed “axo-vascular coupling” (AVC), differed significantly from the NVC that has been studied in grey matter preparations.

The overall pattern of NVC and AVC responses is different. Whereas in cortical slice preparations blood vessels respond rather directly to the rise and fall of neuronal activity (Mishra et al., 2016), the vessel dilation in the ON progresses far beyond the end of the stimulation, reaching a maximal dilation after 20 min and not returning to baseline before 30 min (Figure 3). This “tonic” response contrasts with the “phasic” response of the grey matter (Hall et al., 2014; Mishra et al., 2016). However, this timeframe correlates well with the kinetics of oligodendrocytes which bring the glucose transporter GLUT1 to the cell surface after glutamatergic stimulation within 30 min (Saab et al., 2016). It is also possible that the sustained dilation is caused by the temporal summation of glial signals along the whole nerve and action potential traveling distance, glial cells might remain active long after electrical stimulation, however, this remains to be addressed.

We note that in WM tracts oligodendrocytes and astrocytes are gap junction-coupled (Köhler et al., 2019; Lundgaard et al., 2014; Orthmann-Murphy et al., 2008) similar to endothelial and perivascular cells (Kovacs-Oller et al., 2020; Longden and Nelson, 2016). The pan glial syncytium has been shown to allow an efficient transport of metabolites (Cooper et al., 2020). The vascular syncytium is responsible for the movement of dilatory signals following the vessel order in a decreasing fashion (Longden et al., 2021), at a speed of 27 µm/sec through the endothelial syncytium (Hariharan et al., 2022). With the murine ON stretching for approximately 5 mm, it would take more than 3 min for electrical signals to travel from the most rostral pericyte to the pre-capillary arteriole in the caudal position. Since all ON axons were stimulated at the same time (for 3 min), virtually all pericytes should have received vasodilatation signals. It is thus plausible that there is an accumulation of (metabolic and electrical) signals and intracellular responses that explain the sustained dilation under these experimental conditions.

Although the vascular dilation increase of 7.4%, on average, with all the studied stimulations frequencies, appears small, according to Poiseuille’s law (Pfitzner, 1976), this percentage increase would correspond to a striking 33.1% increase in blood flow *in vivo*, assuming constant blood pressure and dynamic viscosity. The lack of a physiological vascular tone in our, and in previously, studied *ex vivo* systems has a major impact on the temporal delay of NVC, e.g. when compared to the time course of NVC observed by *in vivo* two-photon imaging (Mishra et al., 2016; Rungta et al., 2021).

Interestingly, the oxygen concentration showed a particularly strong effect. In gray matter, a change from 95% to 20% oxygen increases the amplitude of the NVC response (Hall et al., 2014) and attenuates the uptake of PGE_2_, thereby leading to its accumulation and vessel dilation (Gordon et al., 2008). In our WM system, in contrast, the same reduction from 95% to 20% oxygen changed the NVC pathway from a non-prostaglandin mediated mechanism to a PGE_2_-EP_4_ mediated response (Figure 4), and only slightly decreased the amplitude of the NVC response. Although we cannot measure the partial oxygen pressure (pO_2_) in our system, we can extrapolate from published data. A typical 100 µm thick brain slice at 20% oxygen exhibits a pO_2_ of 18.94 mm Hg measured 50 µm below the surface (Gordon et al., 2008). The radius of an ON is ∼150 µm, and the imaged vessels are located 10 µm to 40 µm below the surface. Using the reported values by (Gordon et al., 2008), we calculated that the imaged cells are exposed to a pO_2_ of ∼22 mmHg – 32 mmHg, which is in line with the reported brain pO_2_ values *in vivo* (Lyons et al., 2016; Offenhauser et al., 2005).

We note that the stimulation frequency of ON had no obvious effect on vessel dilation amplitude. In grey matter, for comparison, different stimulated neuronal pathways and brain areas exhibit different changes in cerebral blood flow. Transcallosal fiber stimulation shows a linear correlation between the stimulation frequency and blood flow changes (Enager et al., 2009), possibly due to linear increases in vessel dilation amplitudes. However, in the ON, the vessel dilation does not follow a linear correlation with the stimulation rate in the studied range of 4 Hz – 100 Hz (Figure 3). While the number of ON axons that are unable to repolarize progressively increases when stimulated with frequencies above 15 Hz (Saab et al., 2016; Stys et al., 1991; Trevisiol et al., 2017), they rarely sustain damage, and the ON conduction resumes within 1 min following the end of high-frequency stimulation as shown by the CAP area reaching 95% of the baseline levels after 6 min (Figure 5). The percentage of the stimulation-evoked dilation that is reached at the end of the post-stimulation period is the same for all three frequencies (Figure 3). This would suggest that in an experimental setting with supramaximal stimulation the ON exhibits an all or none response, possibly due to the summation of propagating signals across all pericytes. Experiments with a broader range of stimulation intensities and frequencies between 0.1 Hz and 4 Hz are required to reach a firm conclusion and rule out artifacts of the global stimulation protocol. By using ACh as a positive control at the end of each experiment we could rule out that the prolonged dilation is an artifact of cells that have lost their contractile properties, as ACh needs both endothelial and perivascular cells to be functional.

The most important aspect of our study is the genetic finding that oligodendrocytes play an important role in AVC, presumably by picking up the energetic needs of axons. Compelling evidence suggests that spiking axons release traces of glutamate into the periaxonal space (Doyle et al., 2018; Kukley et al., 2007; Ziskin et al., 2007) that is detected by oligodendrocytes which express NMDA receptors at the inner myelin membrane, facing the internodal axon (Micu et al., 2016; Saab et al., 2016). Here, frequency-dependent axonal glutamate release could be vesicular (for glutamatergic neurons) or non-vesicular in other axons (following sodium transients that revert the direction of glutamate transport), which is still under debate.

Interestingly, in an oligodendrocyte-specific NMDA receptor knock-out mouse, we found that the AVC response is quantitatively impaired, but the time needed to reach maximal dilation is not different than in control nerves (Figure 5). These data would suggest that the oligodendroglial NMDA receptors may not be the only glutamate receptors involved in the AVC response. It is also possible that NMDA receptor expression in OPC contributes to our observation, as Cnp-Cre expression has been observed in a subset of OPC in the ON (Saab et al., 2016).

In grey matter, astrocytes are involved in NVC by relaying synaptic activity to pericytes via PGE_2_-EP_4_ mediated signaling (Gordon et al., 2008; Hall et al., 2014; Mishra et al., 2016). Single-cell RNA sequencing data revealed that the expression of prostaglandin-producing enzymes is restricted to astrocytes (Vanlandewijck et al., 2018; Zhang et al., 2014), which suggests that in WM, similar to grey matter, astrocytes are involved in the AVC response.

Although the link between oligodendroglial NMDA receptor deletion and decreased AVC in the WM remains to be further studied, a key factor could be a second messenger that permeates through the NMDARs. The candidates for this second messenger are calcium and, to lesser extent sodium. It has been shown that in astrocytes, after glutamate exposure, the increase in both sodium and calcium levels induces an increase in GLUT1 receptors (Porras et al., 2008). It is conceivable that the activation of NMDARs in oligodendrocytes also increases the concentration of sodium, which could flow in waves into astrocytes due to both cell types having the same resting sodium concentration (Ballanyi and Kettenmann, 1990; Rose and Chatton, 2016). There is evidence that an increase in sodium can produce a release of calcium from internal stores in cortical astrocytes (Felix et al., 2020; Ziemens et al., 2019) which could be used for PGE_2_ synthesis. Sodium could act as a second messenger responsible for AVC in WM but this remains to be experimentally confirmed (Figure 6).

**Figure 6:**
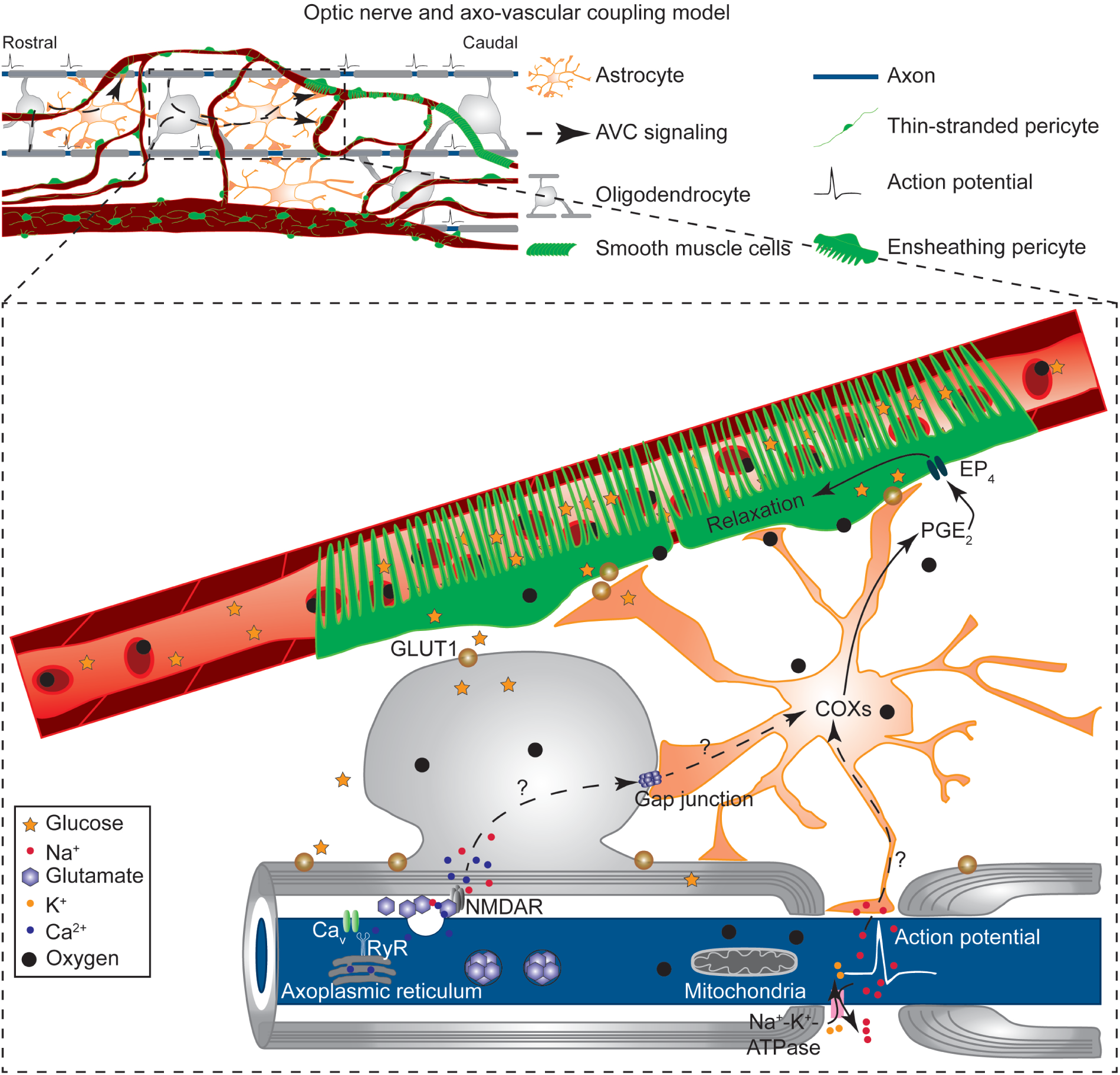
Working model of “axo-vascular” coupling. Axonal activity is detected by oligodendrocytes that initiate a signaling pathway to induce vessel dilation via astrocytes. As the entire ON is stimulated in this set-up, virtually all oligodendrocytes will signal at the same time, inducing an accumulation of vasodilatory signals that remains in the tissue for at least 20 min. Oligodendroglial NMDARs are involved in the AVC, however, other complementary mechanisms can be in place. During the axonal activity, sodium is extruded from the axon at the nodes of Ranvier to maintain the ionic balance. Astrocytes could detect the sodium increase to induce internal calcium release to activate COX enzymes to produce PGE_2_. Additionally, ATP release at the nodes of Ranvier could trigger PGE_2_ release to induce AVC, as shown in grey matter NVC.

In conclusion, we show that the WM AVC is considerably different from the grey matter response. Possibly, this is due to the fundamentally different nature of the tissues and electric signal location. While the grey matter has a confined activation at the synapses, the WM is subject to activation along the entire length of the axons. This suggests that the WM vasculature has the task of supporting a much bigger area. Compensating for a smaller vessel density, it is possible that the vasculature role in WM is to keep pre-capillary arterioles dilated for a longer period of time to allow a sufficient increase in blood flow to match energy needs with an increased oxygen and glucose diffusion from capillaries. This diffusion supports a more efficient axonal ATP production from glial lactate. These characteristics of WM AVC should be considered when studying human WM activation with fMRI experiments, which have been elusive (Gawryluk et al., 2014; Mazerolle et al., 2008).

## Methods

### Ethics statement

All mice used during this work were bred and kept in the animal facility of the Max Planck Institute for Multidisciplinary Sciences. The animals were handled according to the guidelines for the welfare of experimental animals issued by the European Communities Council Directive 2010/63/EU, and the local animal care guidelines of the Max Planck Institute for Multidisciplinary Sciences. Experiments were approved by the institute’s animal welfare officer and the Landesamt für Verbraucherschutz und Lebensmittelsicherheit (LAVES). Animals were housed in standard cages and lived in a 12h/12h light-dark cycle with food and water *ad libitum*.

### Transgenic mice

*Cnp1*-Cre; *Grin1* flox/flox mice described previously (Saab et al., 2016) were crossbred with *Pdgfrβ*-eGFP mice (Gensat.org. Line name: Tg(*Pdgfrb-eGFP*)JN169Gsat/Mmucd). For Figure 3, mice with a *Cnp1*-Cre +/+; *Grin1* flox/flox (no cre expression) genotype were used. For Figure 1, Figure 2, and Figure 4, mice from the reporter line *Pdgfrβ*-eGFP were used. Mice were kept on a mixed C57Bl6 N/J mixed background. Mice were used at 14-16 weeks of age unless otherwise stated.

### ON tissue preparation for light-sheet microscopy

Mice were sacrificed by cervical dislocation. ONs were rapidly removed and fixed with 4% paraformaldehyde (PFA – Merck) diluted in PBS for 16 hours. Fixed ONs were cleared following a modified iDisco protocol (Depp et al., 2021). Tissue was dehydrated in sequential methanol baths (50% 1x, 80% 1x, 100% 2x, 1 hour each) and then bleached with a solution of H2O2:DMSO:methanol (1:1:4 ratio respectively) overnight at 4°C. The tissue was delipidated by incubation in methanol as follows: 100% for 30 min at 4°C, 100% for 30 min at −20°C, 100% overnight at 4°C, and 80% for 2 hours at room temperature. Following the delipidation, samples were gradually rehydrated in descending methanol in PBS series (80% 1x 2 hours, 50% 1x, 0% 1x, 1 hour each) followed by 0.2% Triton X-100 in PBS (2x, 1 hour each). The tissue was permeabilized overnight by incubating the nerves in a solution of PBS/0.2% Triton X-100/ 20% DMSO/ 0.3 M glycine at 37°C. Before the immunolabeling steps, the tissue was blocked in 0.2% Triton X-100, 10% DMSO, and 6% goat serum for 3 days at 37°C and washed with a solution of PBS (0.2%) Tween-20 (10 mg/ml) heparin (5mM) sodium azide (PTwH) (2x, 1 h). ONs were incubated in PTwH + primary antibodies (anti-podocalyxin (1:500, R&D Systems – MAB1556) and anti-eGFP (1:500, Alves labs – GFP-1020) for 14 days at 37°C. Upon completion of primary antibody labeling, tissues were washed in PTwH (6x, 10 min) and stored overnight at 37°C. For secondary antibody labeling, ONs were incubated in PTwH/3% Goat serum + secondary antibody (anti-rat AlexaFluor 555 (1:500, Molecular probes), and anti-chicken AlexaFluor 633 (1:500, Invitrogen)) for 7 days at 37°C. Before clearing, the tissue was embedded in 1.5% w/v Phytagel in water. ONs were washed in PTwH (3x, 10 min each) before dehydration through an ascending concentration of methanol in PBS (20% 1x, 40% 1x, 60% 1x, 80% 1x, 100% 1x, 1 h each at room temperature) and delipidation in a 1:2 mixture of methanol:dichloromethane (overnight at room temperature). Lastly, samples were cleared by immersing them in ethyl cinnamate (Eci, Sigma-Aldrich) until transparent. All incubation steps were carried out at constant medium-speed rotation at the indicated temperatures. Samples were stored at room temperature in Eci until imaging.

### Light-sheet imaging

Cleared samples were imaged using a commercial light-sheet microscope, UltraMicroscope II (LaVision Biotec), equipped with a 2x objective lens, zoom body, and a corrected dipping cap. The tissue was submerged in a sample cuvette containing ethyl cinnamate (ECI). Images were acquired with the mosaic acquisition mode using 5 µm light sheet thickness, 30% sheet width, 0.154 sheet numerical aperture, 4 µm z-step size, 5-step dynamic focus, and 100 msec exposure time. Images were imported into Vision4D v3.2/v3.1 (Arivis) and stitched using the tile sorter configuration. Stitched images were exported and the contrast was adjusted with Fiji (Schindelin et al., 2012).

### Whole-mount ON tissue preparation for confocal microscopy

Mice were anesthetized with a combination of Midazolam (5.0 mg/kg), Medetomidine (0.5 mg/kg), and Fentanyl (0.05 mg/kg) in sterile 0.9% saline solution (MMF) via intraperitoneal injection using 100 µl per 10 g of body weight and injected i.v with 100 µl of 70 kDa Dextran-Texas red (10 mg/ml – Life Technologies, D1830) dissolved in PBS pH 7.4. Mice were decapitated and the ONs were rapidly removed and fixed with 4% paraformaldehyde (PFA – Merck) diluted in PBS for 16 hours. ONs were whole mounted using Aqua-Poly Mount (Polysciences, Inc) in glass slides and imaged as soon as possible.

### Confocal microscopy imaging of fixed nerves

Imaging of the ONs was performed with a Zeiss LSM 880 with Airyscan Fast (Carl Zeiss AG). Images were acquired using a 20x dry objective (Plan-Apochromat 20x/0.80 M27, Carl Zeiss AG).

For figure 1B a z-stack configuration of 89.26 µm was used. The voxel size was 0.30 µm x 0.30 µm x 0.73 µm, the frame size was 5892 px x 2971 px and the image size was 1.75 mm x 880.9 µm. For figure 1C a z-stack configuration of 94.3 µm was used. The voxel size was 0.38 µm x 0.38 µm x 0.73 µm, the frame size was 5900 px x 2970 px and the image size was 2.23 mm x 1.12 mm. For figure 1D a z-stack configuration of 40.13 µm was used. The voxel size was 0.09 µm x 0.09 µm x 2.51 µm, the frame size was 1024 px x 1024 px and the image size was 95.43 µm x 95.43 µm.

### ON preparation and electrophysiological recordings

The acute ON preparation and electrophysiological recordings have been described previously (Trevisiol et al., 2017). Mice were anesthetized with a combination of Midazolam (5.0 mg/kg), Medetomidine (0.5 mg/kg) and Fentanyl (0.05 mg/kg) in sterile 0.9% saline solution (MMF) via intraperitoneal injection using 100 µl per 10 g of body weight. After checking that the animal was in deep anesthesia, mice were injected 100 µl of 70 kDa Dextran-Texas red (10 mg/ml – Life Technologies, D1830) dissolved in PBS pH 7.4 via tail vein.

Mice were decapitated and the ONs were excised as fast as possible without damaging them. One ON was placed on the interface Brain/Tissue Slice (BTS) chamber system (#65-0073; Harvard apparatus), between two custom-made suction electrodes (1.5 mm, #1B150-6, World Precision Instruments) and kept superfused with artificial cerebrospinal fluid (aCSF). The chamber was continuously aerated by humidified carbogen (95% O_2_ + 5% CO_2_) or a mixture of 20% O_2_ + 5% CO_2_ + 75% N_2_. The chamber was kept at 35°C using a feedback-driven temperature controller (TC-10, NPI electronic) connected to a temperature probe (TS-100-S, NPI electronic) inserted directly into the BTS chamber. The second nerve was kept in a solution of aCSF at RT with constant oxygen bubbling until it was time to use it in the BTS chamber.

The stimulating electrode was connected to a battery (Stimulus Isolator A 385, World Precision Instruments) that delivered a supramaximal stimulus of 0.75 mA that evoked the compound action potential (CAP). The recording and reference electrode were connected to the headstage, the signal was pre-amplified 10-fold (EXT-10-2F, NPI electronic) and subsequently 20-fold by a low-noise preamplifier (SR 560, Stanford Research Systems). The amplified signal was low-passed filtered at 30 kHz, defined by the SR 560 preamplifier, and acquired at 20 kHz or 100 kHz by a HEKA EPC9 amplifier (Heka Elektronik). The reference channel was recorded by an aCSF-filled glass capillary, in contact with the bathing aCSF, next to the recording electrode.

Before starting the experiment, the nerves were equilibrated for at least 20 min in the chamber. During the experiments, the nerves were recorded at a baseline stimulation of 0.1 Hz. The stimulation at 25 Hz and 100 Hz was done by burst-like stimulation by applying 100 stimuli at the desired frequency separated by 25 msec during which the CAP was recorded. The stimulation at 4 Hz was done by continuously stimulating the nerve every 250 msec.

### *Ex vivo* vessel imaging

Imaging of the acute excised ONs was performed using a modified up-right confocal laser scanning microscope Zeiss 510 META/NLO (Carl Zeiss AG) equipped with a 63x water immersion objective (Achroplan IR 63x/0.9 W, Carl Zeiss AG).

The pixel size of the images was decided based on the Rayleigh criterion (Waters, 2009). A pixel size (px) of 0.37 µm, a frame size of 256 × 256 px, and an image size of 95.1 × 95.1 µm were used. This resulted in a pixel dwell time of 2.51 µsec and a scan time of 196.61 msec per frame. No averaging was used. Due to the high scattering nature of myelin, an open pinhole was used. The theoretical optical section used was 5.0 µm (2.98 Airy Units). To ensure that the imaged vessel was always being observed in its entirety, a z-stack configuration of 17 slices and an interval between slices of 2.51 µm was used. In total, a z-stack of 40.13 µm was taken. Each z-stack took ∼ 17 secs to acquire. Additionally, there are 3 secs between frames, meaning that a frame was imaged every 20 secs.

The region identification for the experiments was made following two criteria upon visual inspection. 1) the vessel should not be collapsed and 2) the vessel should be covered by a perivascular cell identified visually as a pericyte, based on the “bump-on a log” definition and classified as ensheathing (EP), mesh, or thin-stranded (TSP) based on the morphology (Grant et al., 2019; Mishra et al., 2014).

### Solutions

aCSF containing (in mM) 126 NaCl, 3 KCl, 1.25 NaH_2_PO_4_, 23 NaHCO_3_, 2 MgSO_4_, 2 CaCl_2_, 3.3 Glucose, and 6.7 Sucrose was constantly bubbled with carbogen or the 20% oxygen mixture, depending on the experiment. Inhibitors were diluted first in DMSO (except TTX), following manufacturer instructions, and then in aCSF to the corresponding concentration. U46619 (100 nM – Cayman chemical, 16450), Acetylcholine (100 µM (ACh) – Sigma Aldrich, A6625), Tetrodotoxin (1 µM (TTX) – Alamone labs, T-550), L-161,982 (1 µM – Tocris, 2514), Indomethacin (25 µM – Tocris, 1708). For control experiments of each inhibitor, the same amount of DMSO was used.

## Data analysis

### Pericyte numbers

Pericyte number in the ON was calculated by manually counting the number of pericytes using the cell counter in Fiji (Schindelin et al., 2012). Four random images per ON, of 6 different mice, were used. The maximum intensity projection of each z-stack was used and cell morphology was evaluated for the manual counting.

### Vessel length density

The average vascular length in ONs, corpus callosum, and cortex was calculated using the NeuronJ plugin (Meijering et al., 2004) in Fiji. The volume of each region was calculated by delimiting it by hand. The vessel was then traced using NeuronJ and the result was extrapolated to be in m/mm^3^.

### Vessel diameter

Images for vessel diameter calculation of acutely dissected ONs were acquired using the microscope software Zen 2009 (Carl Zeiss AG). We developed a workflow that allows for a semi-automated quantification of the entire imaged vessel adapting published methods for vessel diameter quantification (McDowell et al., 2021; Mishra et al., 2014). For the analysis, the 17 slices are summed and a median filter with a kernel of 4 px was used to denoise the image. To improve the denoising and reduce the background, a second median filter with a kernel of 30 px was used to create an artificial background that was subtracted from the filtered image.

The summed, filtered image was then registered using different Fiji plugins. StackReg (Thévenaz et al., 1998) was the main plugin used. If this failed to properly register the image, then Image Stabilizer was used (https://imagej.net/plugins/image-stabilizer). In case none of the above plugins work, a manual registration (https://imagej.net/plugins/manual-drift-correction) was performed. Depending on the movement of the nerve, a combination of plugins was used.

The registered image was then segmented using the Li algorithm (Li and Tam, 1998) in Fiji. The segmented image was skeletonized to obtain the center of the vessel. The center of the vessel was used as a guide to drawing regions of interest (ROIs) to calculate the diameter of the vessel. A perpendicular line of 5 px width and 60 px length was used. The intensity profile of the image along that line was calculated and fitted to a Gaussian curve. The diameter of the vessel was then obtained by calculating the Full-Width at Half Maximum (FWHM) of each fitted curve. The perpendicular line sweeps all the ROIs and saves the FWHM of each line, thus calculating the diameter of the vessel at each point during the entire duration of the experiment.

The semi-automated workflow will result in a graph showing the results of the calculation of the FWHM of the vessel measured along perpendicular intensity profiles along the entire vessel. The last 2 min of each part of the experiment (i.e. baseline, preconstriction, dilator, electrical stimulation, or inhibitor) were plotted in a graph showing the FWHM versus the position of the nerve that corresponds to where the intensity profile was measured (Figure 2B). The region of the vessel that will be quantified over time to evaluate vessel dilation, was chosen if it meets two mandatory criteria: 1) the vessel showed a constriction during the application of U46619, a known vasoconstrictor (Mishra et al., 2016), and 2) it dilated during the application of ACh, a known vasodilator (Figure 2B, C). If these two criteria were not met, the quantification of vessel size over time will look quite different (Figure 2C). These two criteria indicate that the vessel, in that region, was capable of constricting and dilating. Not using ACh, or another vasodilator, as positive control could lead to false measurements due to perivascular cells losing their contractile properties over time. In case multiple regions fit these criteria, a visual inspection of the location of the line was performed and if it was clear that they belong to two different pericytes, they are counted as two different measurements, giving an n= 2. This is a common practice in such techniques (Mishra et al., 2014, 2016).

### CAP

The function and overall health of the ON were monitored by using the partial CAP waveform, representing the summation of the simultaneous firing of the axons (Saab et al., 2016; Stys et al., 1991). The area under the curve was calculated by integrating the area that starts approximately 0.2 msec after the stimulus artifact and ends at the lowest point of the second peak, approximately 1.5 msec after the onset of the first peak. The data was relativized to the mean of the baseline (one minute before the start of the stimulation). The data of several ONs were pulled together, averaged, and plotted over time. The variability of the data is shown in those graphs by plotting the standard deviation of the mean.

### Presentation of data

Vessel diameter profiles over time are presented as mean ± SEM. The quantification of the diameter profiles is presented as box plots with the whiskers indicating the maximum and minimum, and the central line of the box represents the median.

Data were checked for normality using the Shapiro-Wilk test (Mishra et al., 2019). If all the groups passed the normality test, then an ordinary one-way ANOVA was performed using the minimum possible multiple comparisons needed or a two-tailed t-test, only if two groups were compared. The specific number of comparisons is indicated in the figure legend or in the text. For the two groups’ comparison, variances were compared using an F-test, if they are different, then the Welch correction was used.

If at least one data set did not pass the normality test, then a Kruskal-Wallis test was performed with multiple comparisons, or only if two groups were compared, then a Mann-Whitney two-tailed test was performed. p-values that are < 0.05 are shown as an exact number unless stated otherwise in the figure legend. For comparisons using the same nerves, paired tests were used. For comparisons using different nerves, unpaired tests were used. All statistics and data representation were performed using GraphPad Prism 9.3.0 (Graphpad Holdings, LLC).

## Acknowledgments

Within the Max Planck Institute for Multidisciplinary Sciences, Göttingen, we thank the light microscopy facility at the City Campus, the mechanical workshop for the construction of the imaging setup, and the technical assistants of the Neurogenetics department for help with genotyping as well as the animal facility for excellent mouse husbandry.

## Author contribution

AR and KAN conceptualized and designed the study. AR performed experiments and analyze data. AT and CRA wrote analyzing scripts. CD and AOS performed tissue clearing and light-sheet microscopy. AK provided mice. IDT and JH provided conceptual input. AR, AT, and KAN wrote the manuscript. All authors revised and approved the final submission.

